# Differential Contributions of CA3 and Entorhinal Cortex Inputs to Ripple Patterns in the Hippocampus Under Cannabidiol

**DOI:** 10.1101/2024.08.06.606645

**Authors:** Adrian Aleman-Zapata, Melisa Maidana Capitan, Anumita Samanta, Pelin Özsezer, Kopal Agarwal, Tugdual Adam, Abdelrahman Rayan, Lisa Genzel

**Author notes:** equal contribution (Adrian Aleman-Zapata, Melisa Maidana Capitan, Abdelrahman Rayan, Lisa Genzel).

## Abstract

Cannabidiol (CBD), increasingly recognized for its potential to treat insomnia, notably extends NonREM sleep phases and modifies sleep-associated ripple dynamics. Utilizing a threshold-based approach, our study differentiated distinct ripple types in rats, clarifying the contributions of intra-hippocampal (CA3) and cortical (mEC) regions to these events. The findings reveal that CBD primarily influences the CA3’s input to the CA1, resulting in an increased occurrence of short ripples predominantly induced by cortical (mEC) activity and a corresponding decrease in long, intra-hippocampal sharp-wave-ripples. This study highlights the critical interplay between the CA3 and entorhinal cortex dynamics in shaping the characteristics of hippocampal ripples under the influence of CBD.

## Introduction

Cannabidiol (CBD) is on the rise as over the counter remedy for many ailments such as chronic pain, anxiety, epilepsy, and sleep disturbances^1–3^. CBD as well as CB1-receptor agonist have been shown to promote sleep in multiple pre-clinical and clinical studies^4,5^. We previously confirmed the sleep-promoting effect of CBD in rats but discovered that CBD also shortens hippocampal ripples during sleep, resulting in disrupted complex but intact simple memories the next day^6^. Specifically, we assessed the ability of rats to recognize object-location patterns over multiple trials and retain this knowledge until testing the following day. We differentiated between simple learning, where the same two locations are consistently reinforced, and complex learning, where one of the two locations remains the same but the other changes from trial to trial. Changes observed solely after simple learning would indicate the consolidation of simple memories, as previously reinforced during training^7–9^. Conversely, changes specific to complex learning likely reflect the consolidation of semantic-like memory, involving the comparison and integration of new and old information^7–9^. Interestingly, while all animals successfully learned the object-location pattern, those treated with CBD failed to retain the complex memory until the next day.

During ripples, cells that were active during the encoding of new memories are reactivated^10–12^, which is known to support memory consolidation during sleep.^13^. Longer ripples contain extended reactivation sequences and may be crucial for consolidating complex experiences that involve more variations and subtle statistical patterns across trials^10–12^.

## Results

To investigate the underlying mechanisms of CBD-attenuated ripples and memory consolidation during sleep, we utilized 32-channel silicone probes to capture the laminar electrophysiological profiles of the hippocampus during urethane-induced sleep-like states (Fig. 1A). By applying the K-means clustering algorithm to a principal component epoch feature space, we identified two states (Fig. S1), splitting the recordings into NREM-like and REM-like periods. CBD induced more sleep state transitions and deepening of the NREM-like state with more, slower, and larger delta waves (Fig. S1). Moreover, CBD slightly lowered the frequency and amplitude of ripples (Fig. S1). These findings, consistent with our prior observations in natural sleep recordings, serve to validate our experimental model by confirming the effects observed in sleep-like states and ripple patterns.^6^.

**Fig. 1.**
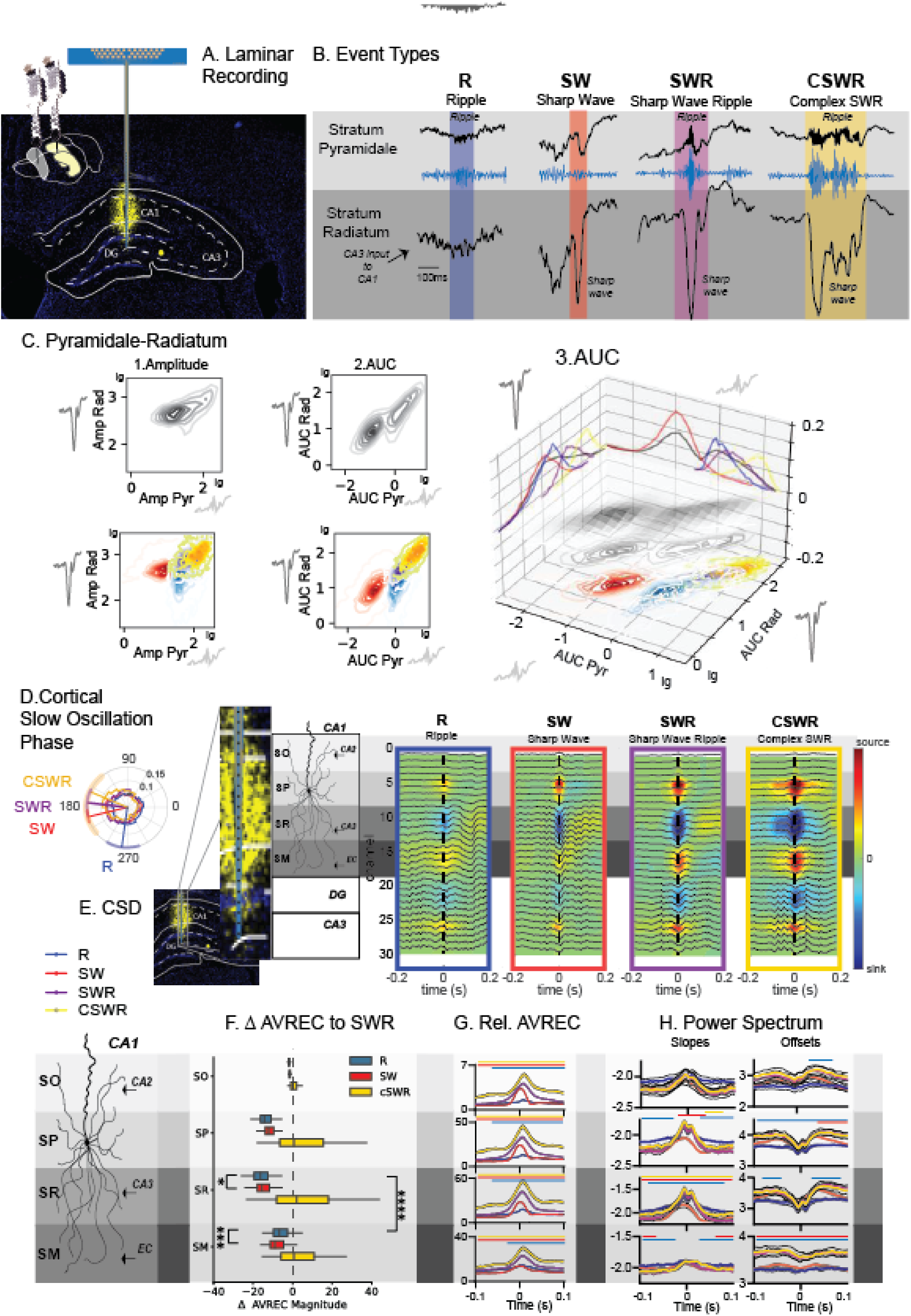
Identifying Ripple Types: A. We performed laminar recordings of CA1 field in the hippocampus B. We applied a threshold-based approach to detect hippocampal ripples in the stratum pyramidale and sharp waves in the stratum radiatum and identified four unique types of events: ripples without sharp waves (R), sharp waves without ripples (SW), sharp wave ripples (SWR) and multiple sharp waves occurring with one long ripple (cSWR). C. Shows amplitude (1) and AUC (2 and 3) of events in the stratum radiatum in relation to the stratum pyramidale. A log-linear relationship is evident only for AUC. D. Events in regard to the slow oscillation phase in the prefrontal cortex. SW, SWR, cSWR occur at the transition of the down to up-state and R in the upstate E. Current Source Density (CSD) maps for each type of event F. Average rectified signal (AveRec) of CSD (ANOVAs for each layer type, time and interaction all p<0.0001) and G. relative AveRec absolute (ANOVA layer F_3,8380_=14930 p<0.0001, type F_3,8380_=90.5 p<0.0001, interaction F_9,8380_=52.97 p<0.0001) and normalized ANOVA (layer F_3,8380_=4.4 p=0.004, type F_3,8380_=99.5 p<0.0001, interaction F_9,8380_=31.8 p<0.0001). Results highlight differential layer contribution to each event type H. Power Spectrum Slopes and Offsets for each layer. For each event type, the power spectrum was computed using a Morlet wavelet transform featuring six cycles over a frequency range of 1 to 200 Hz. We calculated the base-10 logarithms of both the absolute values of the Morlet wavelet power spectrum and the corresponding frequencies. Linear regression analysis of these log-transformed values was then performed to extract the slope and offset values (See Method Section). For F-H shown is mean with SEM (error bars and black lines), lines above the figures indicate p<0.01 Dunnett post-hoc type vs SWR adjusted for multiple comparison. The schematic provides a detailed reconstruction of the CA1 layered architecture. From the outermost to the innermost, the layers depicted are the stratum oriens (SO), stratum pyramidale (SP), stratum radiatum (SR), and stratum lacunosum moleculare (SM), each distinct in its structural and functional contributions to hippocampal circuitry.

The hippocampal ripple, also referred to as sharp wave ripple, is formed by two distinct but temporally correlated events that occur in different layers of the hippocampus^11^. The sharp wave is a large deflection that can be recorded from the stratum radiatum in the CA1 region and is produced by currents in the apical dendrites of CA1 pyramidal neurons triggered by CA3 input^11,14^. This also excites CA1 interneurons, whose interaction with pyramidal neurons induces an oscillation that is detected as a ripple in local field potential recordings of the pyramidal layer^15,16^. Initially, we concentrated on recording channels located in the stratum pyramidale and stratum radiatum of the hippocampal CA1 region, where we independently detected ripples in the former and sharp waves in the latter. In our study, we identified a classical sharp wave-ripple event, characterized by the simultaneous occurrence of both components (SWR, Fig. 1B). Interestingly, we also detected sharp waves without associated ripples (SW) and ripples devoid of sharp waves (R). Additionally, we observed instances of multiple sharp waves coinciding with prolonged, continuous ripples, a phenomenon we have designated as complex sharp wave ripples (cSWR). Given the limitations of our experimental conditions, distinguishing between high-gamma events and ripples without sharp waves was not feasible in the current study. Consequently, in this manuscript, the term *’ripple’* refers to both locally generated ripples and high-gamma events. Analysis of these events revealed that R and SWR generally shared similar durations, whereas SWs were notably shorter and cSWRs exhibited extended durations, as depicted in Fig. S2A. All four events had distinct patterns of power, frequency, amplitude, and area under the curve (AUC) in the stratum pyramidale and stratum radiatum layers as well as differences in the power spectrum in the delta and spindle ranges in the hippocampus and cortex (Fig. S2). Interestingly, the amplitude and AUC in the stratum radiatum displayed diverging patterns across events, with R being smaller and larger than SW, in each feature respectively (Fig.S2A).

The AUC provides a detailed assessment of neural activity by summing up the signal strength and duration, offering a clearer picture of neural behavior over time, unlike amplitude, which only captures the highest point of activity at one moment. This makes AUC especially useful for studying complex or long-lasting neural events, as it accurately reflects both intense and sustained activities. It is also effective for comparing different neural patterns, which are essential for analyzing various neural events, such as ripples. Additionally, the comprehensive approach of AUC reduces the influence of random noise, making it a more reliable measure than solely analyzing the amplitude.

Thus, in correlating the features across both layers, we found that the AUC, unlike the amplitude, exhibited a linear relationship in the log-log representation of the two layers (Fig. 1C). This observation emphasizes the greater efficacy of AUC as a descriptor for capturing the temporal dynamics of distinct events. Given that the stratum radiatum serves as the conduit for CA3 inputs to the CA1 region, and the stratum pyramidale embodies the integration of activity across its dendritic tree, effectively representing the output in response to this input, the linear relationship of the AUC within the log-log space of both layers signifies adherence to a power-law distribution. This observation suggests that the temporal dynamics and the interplay of CA3-EC inputs can generate a multitude of events, with their scale and intensity dependent on the input duration, strength, the resultant degree of neuronal recruitment. The appearance of SW events as a separate cluster from the other events along the log-log relationship revealed that only once the sharp wave component crossed the threshold into the next log magnitude, a ripple was triggered in the SP, resulting in an SWR or cSWR. R events, however, were clustered outside the log-log relationship of the other events.

Interestingly, during the slow cortical oscillation phase, SW, SWR, and cSWR were observed during the transition from the cortical downstate to the upstate. In contrast, Rs were predominantly observed during the upstate, as depicted in Fig. 1D.

To further elucidate the divergent input sources between event types, we calculated the peri-event current source density (CSD) maps along the full recording shank (Fig. 1E). These maps also included the stratum oriens, known for CA2 inputs to CA1, and stratum moleculare, where the medial entorhinal cortex (mEC) innervates CA1. In general, the CSD magnitude increased across events R→SW→SWR→cSWR (Fig. 1F). Upon normalizing the CSD average rectified currents (AVREC) of various events to those observed during SWR, distinct patterns of activity emerged in the stratum pyramidale and stratum radiatum compared to the stratum moleculare, the principal target of entorhinal cortex (EC) inputs (Fig. 1G). Specifically, Rs exhibited a decrease in activity across the stratum oriens, stratum pyramidale, and stratum radiatum, coupled with an increase in stratum moleculare activity. This pattern suggests that, unlike other events, ripples without sharp waves are predominantly driven by medial mEC inputs and, to a lesser extent, by CA3 inputs. The dynamic interplay between CA3 and EC inputs to CA1 is implicated in the genesis of these ripples.

The emergence of ripples is a result of excitation-inhibition balance within the network. The ripple component is initiated by the activation of pyramidal neurons in the CA1^11,17,18^; however, the oscillation is maintained by interneuron activity^19^. While further experimental validation is necessary to establish the spectral slope as a definitive measure of excitatory/inhibitory (E/I) balance within the network, we have explored the spectral and slope profiles of each ripple event type as a descriptive analysis of the reported events. Previous studies suggest that the slope of the power spectrum may indicate the current E/I balance, with flatter slopes suggesting a tilt towards excitation and the power spectrum offsets reflecting overall neural activity ^20^. In our analysis, the R events exhibited flatter baseline slopes and lower baseline offsets (Fig. 1H), aligning with their occurrence during cortical up-states (Fig. 1D). In contrast, the slope increases associated with SW events were shorter and did not reach the magnitudes observed for other events, leading to an earlier decay. Notably, events containing a ripple (R, SWR, cSWR) showed a rapid alternation of power spectrum slopes at the peak of the ripple event, suggesting dynamic interactions between excitatory and inhibitory neurons during ripples^15^.

Our findings demonstrate that a threshold-based approach effectively differentiates various types of hippocampal ripples. Events typically classified as SW, SWR, or cSWR are *<u>largely</u>* determined by the magnitude and nature of CA3 inputs. Conversely, ripples influenced by the mEC typically emerge when mEC inputs coincide with the excitable upstate phase of slow oscillations. Our current study advances the concept that the dynamic interplay between CA3 and EC inputs is critical in generating diverse hippocampal events. Historically, ripples were thought to originate predominantly within the CA3-CA1 circuit. However, our data align with emerging evidence that ripples can also arise from interactions involving both the CA2 region and mEC, highlighting a complex network behavior that supports a variety of ripple types based on the interaction between different hippocampal inputs^21–23^.

Next, we compared all the events detected under the CBD with those from the vehicle condition. Under the influence of the CBD, events exhibited a shorter duration. Notably, the cSWR demonstrated the most pronounced shortening effect (Fig. 2A). Interestingly, there was a distribution shift in the AUC values, particularly for stratum pyramidale, which led to fewer small as well as very large events and thus was again more pronounced for cSWRs (Fig. 2B). In CBD, the log-log relationship between the AUC in stratum pyramidale and stratum radiatum was dampened, with similar stratum radiatum inputs leading to smaller stratum pyramidale responses. This corresponds to CB1 receptors regulating CA3 to CA1 inputs by increasing inhibition at the Schaffer collaterals^24,25^. CSD analysis (Fig. 2D) revealed that CBD treatment resulted in a decreased magnitude of activity in the stratum pyramidale, no significant change in the stratum radiatum, and an increase in the stratum moleculare (Fig. 2D, E). The analysis of the relative average rectified signal (AVREC) demonstrated that the effects of CBD were particularly evident in the stratum pyramidale and stratum radiatum, suggesting a selective impact on CA3 inputs while leaving mEC inputs largely unaffected. Additionally, when comparing these findings to the SWR activity in vehicle-treated animals, a pronounced reduction in input was noted in the stratum radiatum relative to the stratum moleculare. The slopes of the power spectrum were generally less negative and the offsets were lower after CBD intake (Fig. 2G), which was similar to the pattern observed in R events in the vehicle. Finally, CBD intake resulted in more R and fewer cSWR events (Fig. 2H).

**Fig. 2.**
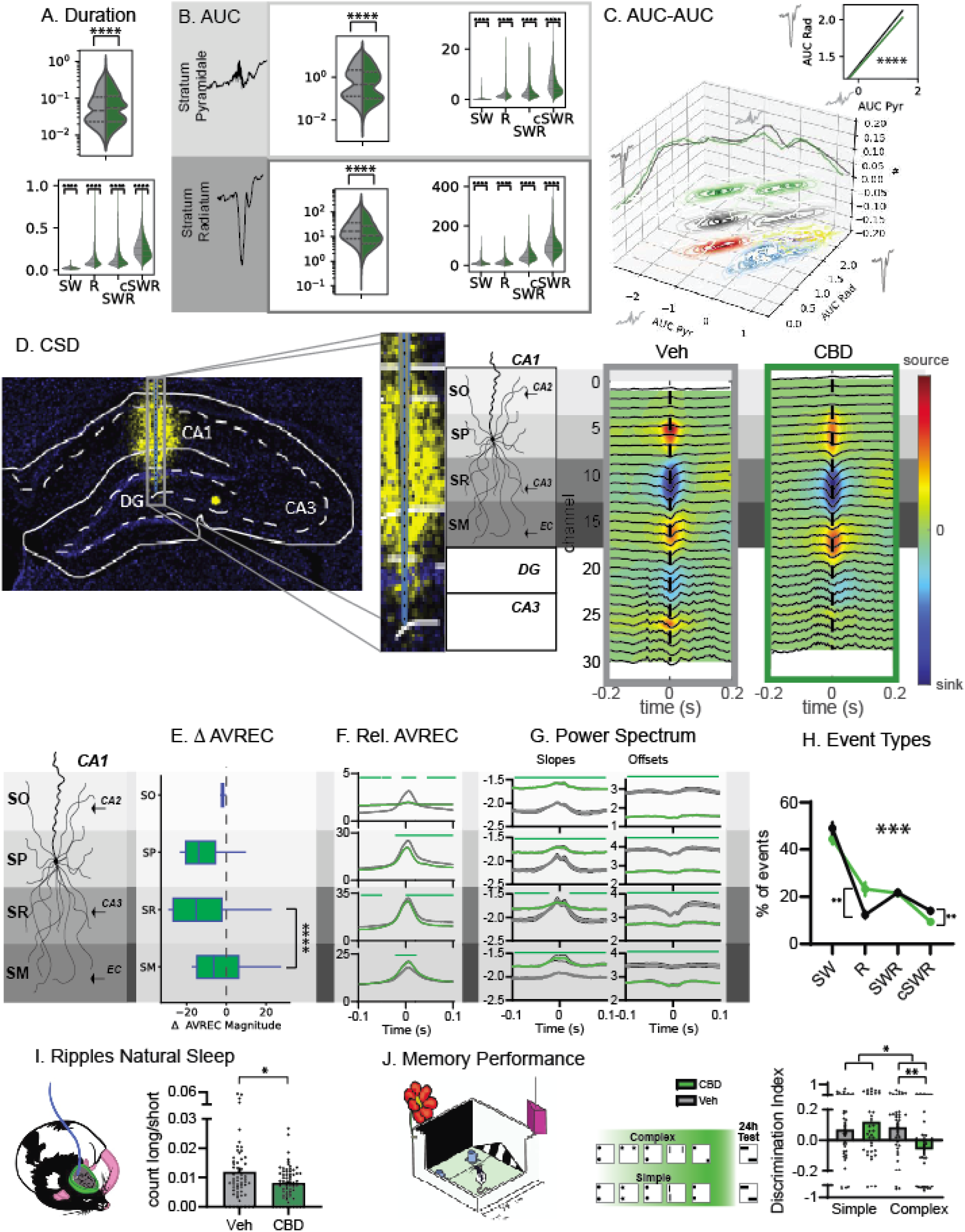
CBD decreases CA3 input to ripples: A. Duration of events shown as probability (top) as well as split for each type (bottom). (Kolm.-Smirnov p<0.0001) B. Area Under the Curve (AUC) for stratum pyramidale and radiatum as probability and split for types. C. Relating AUC from each layer showed a dampening of the log-linear relationship for CBD (Levene p<0.0001). D. CSD maps for vehicle and CBD. E. AveRec showed decreases in SP, no change in SR and increase in SO for CBD CSD (ANOVAs for each layer drug, time and interaction all p<0.0001). F. Relative AveRec across layers with CBD changes evident in SP and SR (ANOVA layer F_3,11328_=1615 p<0.0001, drug F_1,11328_=134.9 p<0.0001, interaction F_3,11328_=48.6 p<0.0001). G. Power spectrum slopes and offsets showed baseline differences for CBD. H. As a result less cSWR but more R were detected in CBD in comparison to Veh (ANOVA drug F_1,560_=0.08 p=0.77, type F_3,560_=124.6 p<0.0001, interaction F_3,560_=6.98 p=0.0001). I. In natural sleep CBD leads to less long in comparison to short ripples (presented here as count long/count short). J. Animals trained on the Object Space Task in complex memory (with one object always in one corner, while the other object moved from trial to trial until the final two trials thus only abstracted gist memory would lead to a positive discrimination index) and simple memory (always same object configuration) showed under CBD worse complex but intact simple memory expression the next day^6^. * p<0.05, **p<0.01***p<0.001. **p<0.01 *p<0.05 E-J Mean with SEM

To confirm our findings in natural sleep, we reanalyzed our previous recordings ^6^ calculating the ratio of short to long occurring ripples and confirmed that CBD intake results in fewer long and more short ripples (Fig. 2I). Next, the Object Space Task (Fig. 2J) was combined with oral administration of CBD. Rats were trained in either the Stable or Overlapping condition, which test simple and cumulative memory respectively. In Stable, in each training trial identical object pairs (each trial different ones) are presented in the same two locations in the exploration box over 5 training trials. At the test (24h after training), two new objects are again used and presented with one on a usual location but the other one placed in one of the other two corners. Neophilic animals will explore the object in the novel location more, independent of whether they just remembered the last experience or all five training trials (leading to positive Discrimination Index, DI). Thus, the Stable condition tests for simple location memory. In contrast, in Overlapping, already during training only one location remains stable, while the second object will be in one of the three other corners. Now, the test trial has the same configuration as the final training trial. Therefore, only if the rats created a cumulative memory, abstracted over multiple training trials, will they show a positive discrimination index at test. All animals underwent both conditions and both treatments, therefore four rounds of training-test, in a cross-over design. Treatment/condition sequences were counterbalanced across rats and object locations. At test, there was a significant condition X treatment interaction (rmANOVA condition F_1,36_=5.6 p=0.023, treatment F_1,36_=1.2 p=0.28, interaction F_1,36_=5.1 p=0.031, Fig. 2J) with memory in the Stable condition intact but no memory expression at 24h in the Overlapping condition (see also ^6^).

## Discussion

In summary, our study has successfully identified four distinct categories of hippocampal events (Fig. 3A), underscoring the fine-tuned dynamics of neural input to the CA1 region. Specifically, our findings highlight the essential role of the interactions within the CA3-EC network in generating different types of ripples. This demonstrates how variations in input dynamics across these regions are fundamental to the diversity of ripple patterns observed in hippocampal activity.

**Fig 3.**
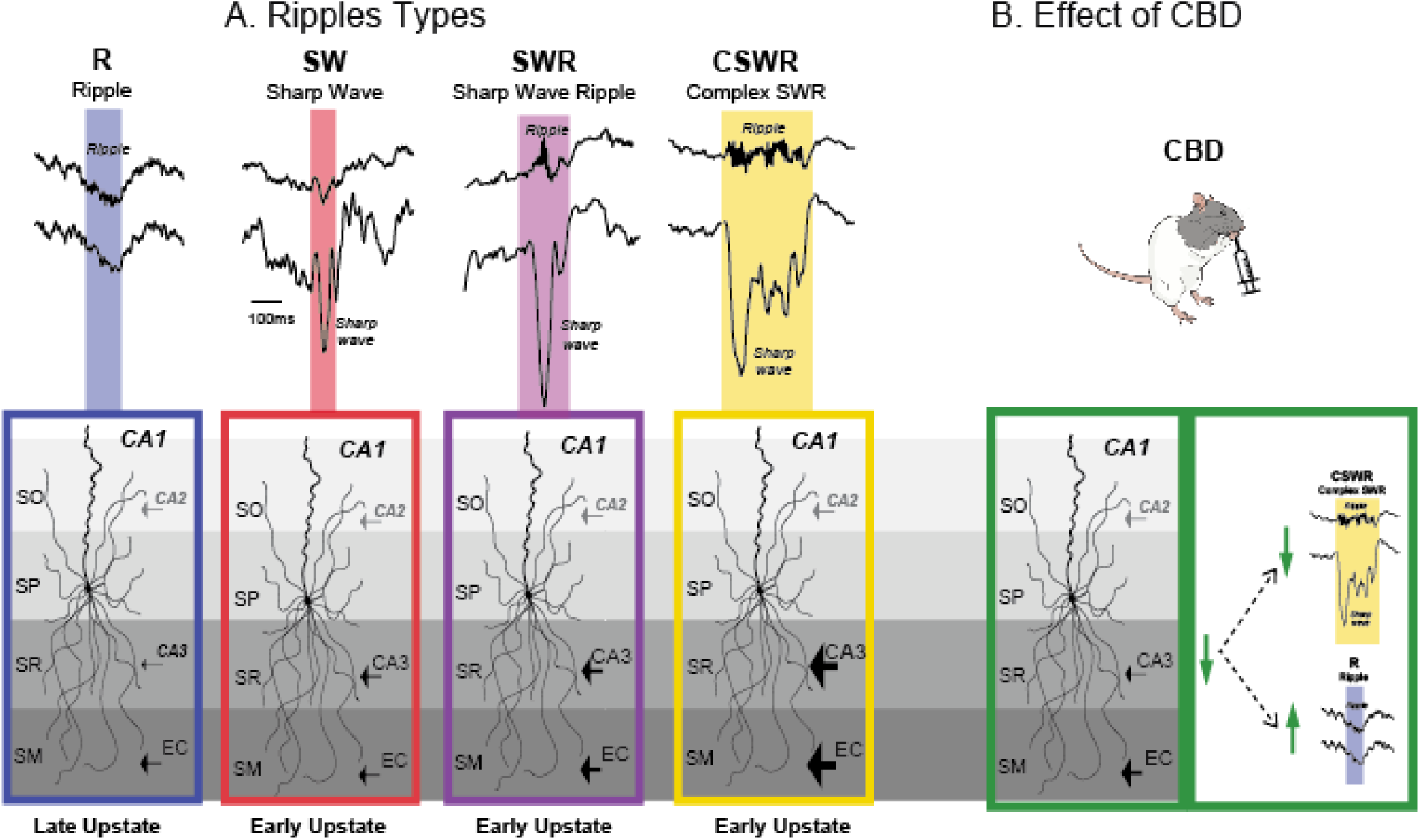
Summary: A. In the present study we applied a threshold-based approach to detect hippocampal ripple and sharp wave oscillations and identified four unique types of events: ripples without sharp waves (R), sharp waves without ripples (SW), sharp wave ripples (SWR) and multiple sharp waves occurring with one long ripple (cSWR). SW, SWR and cSWR shared a common CA3 dominated input pattern, which differed in scale, and events occurred at the transition of the cortical down to upstate. In contrast, R showed a relative higher mEC input and these events occurred in the cortical upstate. B. CBD intake selectively inhibited CA3 input to CA1, which led to the occurrence of fewer cSWR and more R.

In our study, ripples devoid of sharp waves were primarily associated with stronger inputs from the medial entorhinal cortex and typically occurred during the cortical upstate. In contrast, other identified ripple categories were predominantly influenced by stronger inputs from CA3, with stronger inputs generally correlating with longer ripple durations. These CA3-mediated events occurred primarily during transitions from the cortical downstate to upstate.

Notably, the administration of CBD led to a significant reduction in CA3 input, which in turn promoted an increase in the occurrence of ripples lacking sharp waves and a decrease in prolonged, complex CA3-driven sharp wave ripples. This reduction in CA3 input also accounts for the observed disruption in the temporal coordination of cell assemblies within CA1 following activation of CB1 receptors ^6^.

Our previous study has demonstrated that CBD shortens ripples during natural sleep, impairing complex memory while sparing simple memory on the subsequent test day^6^. This study further explores the mechanistic basis of CBD’s selective impact by revealing that it predominantly affects CA3-to-CA1 synaptic inputs, while inputs from the mEC are less affected. Our findings suggest that this specific modulation is governed by critical dynamics within the CA3-EC network, as indicated by our area under the curve (AUC) analysis, which results in a diverse range of ripple activities. The differential impact of CBD on CA3 and mEC inputs has significant implications for memory consolidation processes. Specifically, ripples originating primarily from mEC inputs to CA1 appear to be sufficient for the consolidation of simple memories, which are typically reinforced by repeated experiences during training. Conversely, ripples primarily driven by CA3 inputs, which are longer in duration, are crucial when diverse experiences need to be synthesized to extract subtle statistics over time, as required in complex memory scenarios in our Object Space Task^6,7,9^.

Numerous studies have employed techniques like self-organizing maps and topological data analysis to differentiate ripple types in the pyramidal layer unsupervisedly^26–29^. One notable approach indicated that ripple subtypes could be distinguished by their association, or lack thereof, with sharp waves^28^. Our study demonstrates that a simple threshold-based approach is effective in distinguishing between various ripple types and their associated inputs, provided that events are independently detected across different layers. Furthermore, we have introduced novel categories of hippocampal events, illustrating that CA3 inputs can independently generate sharp waves without necessarily triggering ripples. Additionally, our research has identified instances where ripples, driven primarily by the mEC, occur independently of sharp waves. These findings underscore the fine dynamics within the hippocampal network and suggest a more complex interplay of neural inputs than previously understood.

## Methods

### Study design

A total of 30 rats were used in this experiment. All rats were first extensively handled for multiple days until they experienced minimal stress working with the experimenters (see handling videos on www.genzellab.com). The rats were first habituated to the oral feeding regime and the behaviour training box for a week. In the following week, they were orally administered with cannabidiol (CBD) or vehicle (VEH) (counterbalanced within a single animal) and trained in the stable and overlapping condition of the Object Space task (all conditions counterbalanced within each animal) in smaller sub batches, results published in^6^. Finally at the end of behaviour experiments, these rats were further used for acute electrophysiological recordings. They were orally administered with either CBD or VEH followed by terminal urethane anaesthesia and then implanted with silicon probes in the medial prefrontal cortex (mPFC) and CA1 region of dorsal hippocampus (HPC). The brain activity under anaesthesia was recorded for 6 hours after which the animals were perfused and the brains extracted for histological placement experiments.

### Animals

Six-eight week old male Lister Hooded rats weighing between 250-300 g at the start of the experiment (Charles Rivers, Germany) were used in this study. They were pair-housed in conventional eurostandard type IV cages (Techniplastic, UK) in a temperature-controlled (20 + 2 °C) room following a 12 h light/dark cycle with water and food provided *ad libitum*. A total of 30 rats were used in this experiment: The behavioural experiments (previously published^6^) and electrophysiological recordings were performed during the light period (between 9:00-18:00). All animal procedures were approved by the Central Commissie Dierproeven (CCD) and conducted according to the Experiments on Animals Act (protocol codes, 2016-014-024).

### Drugs

(-)-Trans-Cannabidiol (CBD, > 98%) was obtained from CBDepot.eu for all experiments. Rats were treated with either CBD (120 mg/kg in 300 ul flavoured olive oil, p.o.) or vehicle (300 ul flavoured olive oil, p.o.). Different flavoring agents, namely, vanilla, cinnamon, star anise, and clove, were used to make multiple flavors of olive oil, which were then used to mask the taste of CBD, so that the rats would always be naïve to what they were fed. The use of all flavors was counterbalanced, such that each rat received all flavors for both CBD and VEH. CBD solution was freshly prepared in one of the flavored oils prior to oral administration. To prepare the solution, the amount of CBD to be administered was determined based on the weight of the rat, and the compound was weighed accordingly. The flavored oil was heated to a temperature of 50-60°C and then the CBD compound was slowly dissolved. This process took a few minutes until a clear solution was obtained, which was indistinguishable from vehicle oil. Individual syringes with 300 µL of CBD or VEH were prepared for each rat. The experimenter who performed the oral administration was blinded to the treatment each rat received, and the drug was always administered an hour before the start of behavior training or anesthesia. Previous studies have shown oral administration of CBD was effective in crossing the blood brain barrier and plasma concentrations were shown to reach their maximum concentration around 4-6 hours after oral administration ^30,31^.

### Acute recordings

At the end of the behavioral recordings, 30 rats were used for acute electrophysiological recordings to monitor the effects of CBD on sleep-like states under anesthesia using silicon probes. The rats received an oral administration of CBD or VEH (120 mg/kg) at ∼10 am, and 30 min later received an i.p. injection of urethane anesthesia (1.4 g/kg). After injection, we waited for the next 30 min for the anesthesia to set in and started with surgery at ∼11 am. The aim was to start surgery an hour after the oral administration of CBD/VEH.

### Stererotaxic surgery

Shortly before the start of surgery, all rats received a subcutaneous injection of carprofen (5 mg/kg) to serve as an analgesic. On setting the rat into the stererotax, they further received an s.c. injection of a mixture of 4 mg/kg lidocaine and 1 mg/kg bupivacaine in 0.9% NaCl physiological serum locally at the skin surface above the skull as a local analgesic. Lastly, they also received 2 ml of 0.9% NaCl physiological serum subcutaneous injection at the start and end of surgery. The target areas for the recordings were medial prefrontal cortex (mPFC) and hippocampus (HPC) with the following coordinates – AP = 3.5 mm, ML = 0.5 mm and DV = 2.6 mm (from brain surface) for PFC and AP = -3.2 mm, ML = 2 mm and DV = 4.3 mm (from brain surface) for HPC. All coordinates were calculated with respect to standard bregma and lambda coordinates (Büttner-Ennever, 1997). Two craniotomies (2 × 1 mm and 1 × 1 mm for the PFC and HPC, respectively) were drilled above the target areas in the right hemisphere and a hole in the left cerebellum was drilled for the ground screw. Finally, once the craniotomies were cleaned and the dura mater was clearly visible, silicon probes were placed in position and lowered slowly to both targets. Probe tips were coated with DIL stain (catalog no. D282) before lowering to facilitate better visualization of the electrode damage in the target region during histology.

### Recordings

The surgical setup was enclosed in a Faraday cage to prevent electrical noise in the recordings. Both silicon probes (32 channels in each probe) were connected to an OpenEphys acquisition box (Siegle et al., 2017). The signal was visualized using an OpenEphys GUI and recorded at a sampling rate of 30kHz. Signals from both brain regions were recorded for 6-7 hours and were checked every hour to determine whether the craniotomies were hydrated and the temperature of the heatpad remained constant. At the end of the recordings, the animals were sacrificed via transcardial perfusion, and their brains were extracted for histological analyses.

#### Histology

##### Brain processing

Rats from both acute and chronic recordings were sacrificed via transcardial perfusion at the end of data collection. For acute recordings, the rats were under deep urethane anesthesia at the time of perfusion. For chronic recordings, rats were overdosed with 150 mg/kg sodium pentobarbitol/i. p. prior to perfusion. The rats were perfused first with 100 ml of 0.1 M phosphate-buffered saline (PBS) pH 7.4, followed by 250 ml of 4% paraformaldehyde (PFA) made in 0.1 M PBS. After extracting the brains, they were stored overnight in PFA at 4°C. The brains were rinsed in 0.1 M PBS the next day (3 × 10 min) and then kept in a solution of 30% sucrose and 0.02% NaN_3_ in PBS for cryoprotection. Once the brains sank to the bottom of the vial, they were frozen in dry ice and stored for a long term in -80°C freezer. For further processing, the brains were sectioned in a cryostat (SLEE Medical, Germany) and 50 micron coronal sections of target regions were obtained and collected in 48-well plates containing 0.02 % NaN_3_ in PBS and stored at 4°C. Additionally, the brain sections from the acute datasets were stored in a dark place because the target regions contained fluorescent DIL dye from the probe tips.

##### DIL staining

To visualize the probe location in the brains of the acute datasets, target sections from the prelimbic cortex (PFC) and hippocampus (HPC) were first sequentially mounted (in increasing AP coordinates) on microscopic slides, incubated with DAPI (Abcam, ab104139) for 5 min, and then coated with a coverslip. They were later visualized under the LEICA Thunder Wide-Field Fluorescence microscope, and images were taken at 10X magnification.

#### Local field potential analysis

Here, we describe the methods used to process and analyze the local field potential data acquired during acute recordings. From the 30 recorded rats, 12 rats were excluded of the study (2 rats died during the experiment, 6 rats were missing a brain recording, the pilot rat was removed, 1 rat had corrupted recordings, 1 animal had abnormally short recordings and 1 animal showed abnormal reaction to the drug and abnormal brain recordings). In the end, a total of 18 animals were kept for the data analysis (10 CBD, 8 vehicles). The scripts used to implement the analyses can be found in the following github repository https://github.com/genzellab/cbd. A downsampled version of the LFP data can be found at the OSF website https://osf.io/htmqu/.

##### Scoring of sleep-like states

An automatic state classifier was designed to detect NonREM-like and REM-like epochs based on the spectral and amplitude features of cortical and hippocampal recordings^32^. The spectral power of selected hippocampus and prefrontal cortex channels was estimated from 0 to 100 Hz, with a 0.5 Hz step, using a multitaper filter with a time-bandwidth product of 4 in 10-second-long epochs. Epochs with artifacts were detected using the *isoutlier* MATLAB function on the absolute value of epoch amplitudes. The artifacts were removed and blanked using the signal mean. Principal component analysis (PCA) was employed to identify features of the epochs that characterized the two sleep-like states and explained most of the variance in the data. The following features were used as inputs for PCA: Slow oscillation power (0.1-1 Hz), delta power (1-3 Hz), theta power (3-6 Hz), low beta power (10-20 Hz), low gamma power (30-45 Hz), high gamma power (55-80 Hz), ripple power (90-300 Hz), theta-slow oscillation ratio, and amplitude of the 10-second-long epoch. The frequency ranges were intentionally lower than those reported in the literature, as urethane anesthesia has been shown to slow down brain activity^33^. The first and second principal components were retained, given that they explained most of the variance (∼90%), and a k-means clustering algorithm with two partitions was computed in the PC1-PC2 state space to identify epochs that belonged to the NonREM-like and REM-like state clusters. The first PCA component (PC1) contained high weights for features such as the epoch amplitude and slow oscillation power in both brain areas, which are expected to be high in the NonREM stage during natural sleep. Therefore, the cluster of epochs that contained higher values for PC1 was labeled as NonREM-like sleep, whereas the remaining cluster was labeled as REM-like sleep. For a more extensive description and validation please consult Adam ^32^.

##### Sleep-like state architecture analysis

The percentages of time for the NonREM-like and REM-like states with respect to the total sleep-like time were calculated for groups of consecutive bins: Bins 1-3, bins 4-6 and bins 7-9. Each bin had a duration of 45 minutes. Similarly, the number of transitions between the sleep-like states was counted per bin triplet. For state transitions and their percentages of time, values were computed per rat and averaged across rats that received the same treatment. The mean per bin triplet and its corresponding standard error of the mean were plotted to identify potentially significant changes between treatments. Moreover, the durations of individual sleep-like state bouts were pooled across rats that received the same treatment to identify the distribution of durations for NonREM-like and REM-like states. For more information about sleep-like state architecture analysis, please see Adam ^32^.

##### Preprocessing of recordings

The best channels for the hippocampal stratum pyramidale and stratum radiatum were selected for each rat. The selection criteria consisted of selecting the channels in the vicinity of the layers in which ripples and sharp waves were more prominent. In addition, the superficial and deep channels of the prefrontal cortex were selected. The channels were downsampled from 30 kHz to 600 Hz for use in the local field potential analysis. A 3rd order Butterworth low pass zero-phase filter with a cut-off of 300 Hz was used to prevent aliasing before downsampling the signal. The first 15 min of the recordings were discarded to control for brain signal instability after probe implantation. Artifacts were detected and removed by applying thresholds to the sum of the absolute values of the unfiltered recordings of the selected hippocampal and prefrontal channels. When an artifact was detected, a buildup of 0.5 seconds prior to the artifact and a washout period of 3.5 seconds after the artifact were removed and replaced with the mean value of the artifact-free signal. Before “blanking” the artifacts, we bandpass-filtered the channels which would later be needed for the detection of oscillatory events. This was done to prevent the addition of spurious high-frequency events that occur owing to filtering signal discontinuities after artifact blanking. To study the temporal effect of CBD treatment, recordings from different rats were temporally aligned with respect to CBD feeding time, which was at the same time of the day for all rats. Given that the duration of surgery varied per rat, the starting time of the recordings differed across rats, which made this alignment step important. To account for this variability, we identified the rat whose recording started earlier during the day and temporally aligned the recordings from the remaining rats. This was performed by adding the corresponding amount of NaNs to the start of the recordings of each rat. Similarly, the rat whose recording ended at the latest time of the day was identified, and NaN values were added to the end of the remaining rats’ recordings. The result of this procedure was an aligned matrix per brain area and per layer, in which each row consisted of a recording per rat of a given treatment and the columns corresponded to the total number of temporal samples. We later split this aligned matrix into 45 min bins to study the temporal effects of CBD. A similar aligned matrix was created for the sleep-like state data, which consisted of numerical labels encoding the sleep-like stage for each 10-second epoch. The matrix was upsampled to a sampling rate of 600 Hz to obtain the same dimensions as those of the brain recordings.

##### Detection of hippocampal oscillations

As mentioned in the previous section, temporally aligned matrices were computed for each brain area layer and treatment. To detect hippocampal events, we used aligned matrices that contained bandpassed channels of the hippocampal stratum pyramidale and stratum radiatum. For the stratum pyramidale, the channels were bandpassed in the range 90-200 Hz. For the stratum radiatum, channels were bandpassed in the 2-20 Hz range. Both the raw and filtered versions of the hippocampal channels were displayed in a graphical user interface (GUI), which displayed an amplitude threshold for ripples on the filtered stratum pyramidale channel and an amplitude threshold for sharp waves on the filtered stratum radiatum channel. The GUI presented putative detections based on a threshold of 5 standard deviations with respect to the mean of the filtered signals. When necessary, the thresholds were adjusted for each rat after visually verifying the correct detection of ripples and sharp waves. However, most thresholds remained close to 5 standard deviations above the mean. Once the thresholds were determined, we ran the ripple detection by thresholding voltage peaks that lasted for a minimum duration of 50 ms above the threshold. The start and end of the ripple were determined as half the value of the detection threshold. A closeness threshold of 80 ms was used to count ripples occurring within the proximity of each other as a single event. Sharp waves were detected by finding the voltage troughs below a set threshold. The timestamps in which the troughs crossed the threshold were used to define the start and end of the sharp wave. The sharp wave (negative) peak was the timestamp with the lowest amplitude value between the start and end of the sharp wave. Once events had been detected *<u>independently</u>* in both the stratum pyramidale and stratum radiatum, we looked for their temporal overlap by detecting two possible and non-mutually exclusive cases: 1) The end of the sharp wave was between the start and the end of the ripple, 2) the start of the sharp wave was between the start and end of the ripple. We counted the number of overlaps per ripple to classify the detections as single sharp waves (SW), single ripples (R), sharp wave ripples (SWR) and complex sharp wave ripples (cSWR). Events were classified as SW or R when no overlap was observed. Ripples that overlapped with one sharp wave were classified as the SWR. Ripples that overlapped multiple sharp waves were classified as cSWR. Finally, information on all detections was saved in rows of a table that included the event type; the start, peak, and end timestamps of the event; the sleep-like state during which it was detected; and the 45-minute long bin index in which it was detected. For all the following analyses, we only used the detections that occurred during the NonREM-like stage.

##### Detection of cortical delta waves

The FindDeltaWaves function from the Freely Moving Animal (FMA) toolbox ^34^ was adapted to detect delta waves (1-6 Hz) with a duration between 150 and 550 ms. The peak and trough amplitude thresholds were adapted for each rat after visual validation of the detections. The number of delta waves was determined for each 45 minute-long bin.

##### Slow oscillation phase analysis

Using a 3^rd^ order Butterworth filter the aligned PFC signal (600 Hz sampling rate) was filtered to 0.5-4 Hz and NonREM-like bouts were then extracted to form a concatenated NonREM-like signal. The Hilbert transform of this signal was computed to determine the phase of the slow-wave oscillations in the range of 0°–360°. The concatenated phase signal was then split back into its corresponding sleep state bouts, and the peaks per hippocampal event type were used to determine their corresponding phases. This process was performed for all hippocampal event types in all the rats.

##### Ripple rate at NREM-like

To compare the results of this study with those of our previous research on natural sleep (ref), we conducted the following analysis: First, we calculated the ripple rates at the start and end of NonREM-like bouts. Only bouts lasting longer than 15 minutes were included in this analysis. We isolated the first and last 10% of the duration of each selected bout. Within these segments, we counted the ripple peak timestamps to determine the number of ripples occurring at both the start and end of each bout. The rates for the beginning and end of the bouts were then calculated by dividing the ripple counts by the duration of their respective 10% segments (see Supplementary Figure 1). This method specifically provided the ripple rates for the start and end periods of NonREM-like bouts, distinguishing them from individual event rates. However, it is important to note that the ripple rates reported within the manuscript were computed per 45 bins and across the entire sleep duration.

##### Oscillations characteristics

The traces of each detected event (ripple and delta waves) were extracted using the start and end timestamps obtained from the detectors. Traces of the events were filtered through their corresponding detection frequency bands. Characteristics such as amplitude and mean frequency were calculated for these filtered events using built-in and custom MATLAB functions. The amplitude of the events was calculated by computing the envelope of the filtered trace using Hilbert transform. The absolute value of the result was obtained, and its maximum value was found. The mean frequency of the filtered traces was computed using the MATLAB meanfreq function.

##### Event features in the Radiatum and Pyramidal layers

Several features were computed for simultaneous activity in the stratum pyramidale and radiatum layers during detected hippocampal events. Features were computed from the filtered stratum pyramidale (90-200 Hz) and stratum radiatum (2-20 Hz). All features were calculated using the start and end timestamps of the detections as the temporal limits to compute the features from. In the cases of single sharp waves and ripples, the limits were defined by either the sharp wave or ripple duration, and features were also computed from the simultaneous activity in the corresponding eventless channel. In the cases of the SWR and cSWR, the features were computed in the window given by the start and end of the ripple. This allowed us to capture the activity of multiple sharp waves present during the cSWR. In addition to the amplitude and mean frequency of the events described in the previous section, other features such as the event duration and area under the curve (AUC) were computed. AUC was computed by applying trapezoidal numerical integration to the absolute value of the filtered event with the following formula:

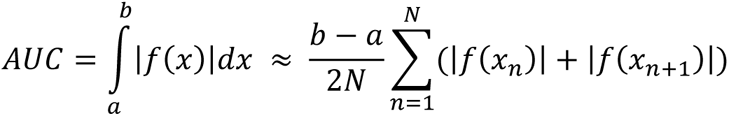

Where *a* and *b* are the start and end timestamps of the hippocampal event, *f*(*x*) is the trace of the filtered event, and *N* is the number of equally spaced samples between *a* and *b*. The absolute value of *f*(*x*) was used to account for negative values. Note that this AUC differs from the amplitude-normalized AUC. By refraining from normalizing the absolute value of the event’s amplitude, we ensured that the AUC value would be influenced by both the trace amplitude and event duration, thereby providing a more accurate estimate of the magnitude of the event. This was particularly useful for describing the magnitude of activity in the radiatum layer during cSWR, which would not have been possible using only the amplitude or duration. Power was computed as the squared Euclidean norm (i.e., vector magnitude) of a ripple waveform divided by its length in samples, following the definition of the energy of a signal over its duration. The number of peaks on a bandpassed waveform was computed using the MATLAB function findpeaks(), which found local maxima defined as data samples larger than its two neighboring samples without a minimum prominence. The sharp-waves frequency was computed differently than the *mean frequency* method employed for ripples. A spectrogram from 1 to 200 Hz was computed using a Morlet wavelet transform on a 6 second-long stratum radiatum trace centered on the sharp-wave peak. The power values between the start and end of the sharp wave were summed over time, resulting in an array with the same length as the number of frequencies. The absolute value of the array was computed and normalized by dividing by its sum. The dot product of the normalized array and frequencies from 1 to 200 Hz were computed. The resulting value was used as the frequency of the sharp waves. Contour plots indicating the density of events in the AUC-AUC plot were computed using a bivariate kernel density estimator implemented with the *gkde2* MATLAB function^35^. To visualize the density as contours, the probability density function was partitioned into three levels of the same density proportion. The probability density function, in which hippocampal events were not grouped by their types, was partitioned into six levels to improve visualization. For such an analysis, we took the same number of random events per type and z-scored the resulting probability density function from the mixed events. The linear fit of the data in the log-log plots was computed using the *polyfit* MATLAB function with an order of 1. Data were first converted to a log10 scale. Pairwise statistical tests between the slopes of the fit lines were implemented by comparing their correlation coefficients using the *corr_rtest* MATLAB function^36^. The test consisted of a z-test of the difference in Fisher’s z-transformed correlations divided by the standard error of the difference.

##### Data preprocessing for power spectra

Artifact rejection was performed by applying thresholds visually determined for each rat. When an artifact was detected, the signal 0.5 seconds before and 3.33 seconds after the artifact was removed to replace the artifact itself and its build-up with the mean of the artifact-less whole signal of each rat^37^. Each event type (i.e., ripple, SW, SWR, cSWR) was extracted as a 6-second long time window of the raw signal using the event’s peak timestamps of each rat event for further processing of the events and conducting current source density analysis. In the case of the SWR and cSWR, the peak corresponds to the event detected in the pyramidal layer. The extracted event windows contained simultaneous activity in each channel (i.e., PFC shallow, PFC deep, HPC stratum pyramidale, stratum radiatum) for all treatments (i.e., Vehicle and CBD) and event types (i.e., random NonREM-like baseline, ripple, SW, SWR, cSWR). All events were sorted with respect to their amplitudes and two thousand events closest to the median for each event type were extracted. Additional artifact rejection was performed on the data of these 2000 events for the PFC shallow and deep PFC. They were band-pass filtered at 100-300 Hz to visually detect the thresholds and remove artifacts. Note that the filter was only used to detect artifacts and the raw signals used in the analysis were not filtered.

##### Power spectra of Stratum Pyramidale and Cortex

Power spectral density (PSD) was computed for acute recordings (i.e., PFC shallow channel and HPC stratum pyramidale). They were computed for both Vehicle and CBD treatments and for each event type (i.e., random NonREM-like baseline, ripple, SW, SWR, cSWR). Baselines were randomly selected NonREM-like periods of 4 s. First, the Fieldtrip toolbox was used to compute PSD^38^. PSD was computed from 0 to 100 Hz in steps of 0.25 Hz and with a length of 4 s. Line noise was removed at 50 Hz using a notch filter. A Hanning window taper was used with a length of 4 s and a 0.25-second overlap to minimize the effect of edge artifacts. To generate the power spectra, PSD values were computed as logarithms with a base of 10.

##### Data Processing for Current Source Density (CSD)

Only a subset of animals had the recording electrodes placed to allow for CSD analysis (n=3 veh, n=2 CBD). The recordings had a sampling rate of 30 kHz. They were downsampled to 600 Hz and arranged in a particular folder structure. Each channel was renamed based on its depth. There were 32 channels in total and the most shallow channel was renamed as ‘1’ and the deepest as ‘32.’ Artifact rejection and aligning were performed in the same manner as for other analyses of acute recordings.

Each event type (i.e., ripple, SW, SWR, cSWR) was extracted with a time window of 6 s centered at the event peak from the raw signal using the timestamps of each event (type) of each rat for each channel. Event types were saved in one file for each rat. The data were converted into cell arrays, where each cell had a size of 32-by-3601 (i.e., 32 being the channel number and 3601 being the number of samples corresponding to 6 s).

#### Current Source Density (CSD) analysis

Neural cellular activity is characterized by current flow through the cell membrane of neurons. This current flow arises as the product of accompanying changes in the biophysical distribution of ions within the extracellular space, giving rise to sinks (inward current flow) and sources (outward current flow). In the current study, we inferred the CSD signal by computing the second spatial derivative of the LFP signal, as previously described^39,40^:

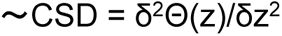

where Θ is the local field potential signal and z is the physical channel on the recording probe representing the spatial coordinate of the channel within the recording space. The CSD was computed for all event types and aligned either to the peak of different event types.

#### Average Rectified Signal (AVREC)

Utilizing individual event CSD profiles, we processed the CSD data by rectifying and then averaging the waveforms across each channel that constitutes the laminar CSD profile, a process denoted as AVREC^39,41^. The resultant AVREC waveform serves as an indicator of the temporal dynamics of each event type, offering a quantifiable measure of the aggregate strength of transmembrane current flow throughout the observed period.

The mean AVREC across all channels contributing to the laminar profile for each oscillatory event was calculated in accordance with the methodology previously outlined by El-Tabbal, et al. ^39^:

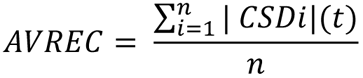

To assess relative changes in input across different layers, we calculated delta AVREC values with respect to SWR events. This involved computing the mean AVREC value for all SWR events for each layer. Subsequently, for each ripple type event, we subtracted these mean values from the corresponding AVREC values of the SWR events, thereby quantifying the relative change in input for each layer compared to the SWR.

#### Slope Analysis of Current Source Density Signal

Conventional slope analysis of the local field potential (LFP) signal power spectrum is used to explore the potential role of the excitation-inhibition balance in various cognitive phenomena^42^. However, the interpretation of LFP signals is challenging because of their origin in neural activity generated by transmembrane currents in adjacent tissues and the confounding effects of volume conduction^43^. In contrast, the current source density (CSD) analysis of LFP recordings delineates regions of positive and negative CSD as sources and sinks, respectively, offering a more direct association with neural activity^43,44^. Biophysically, the CSD more accurately reflects the underlying neural dynamics, establishing it as our primary variable of interest. Consequently, we applied slope analysis to the CSD signal rather than to the LFP to capitalize on its enhanced specificity to neural activity.

In the present study, we computed the power spectrum for the slope analysis of each event type using a Morlet wavelet transform (six cycles, across a frequency range of 1 to 200 Hz) employing the SciPy signal processing toolbox. We then calculated the base-10 logarithm of the absolute values of the Morlet wavelet power spectrum as well as the base-10 logarithm of the corresponding frequencies. Subsequently, these log-transformed values were subjected to linear regression analysis utilizing the linear regression function from the scikit-learn library, from which we extracted the slope and offset values. The analysis window was defined as 250 ms, before and after the event peak.

## Acknowledgments

We would like to thank Victor Canales Lima, Swatantra Dhara, Linda Joseph Tomy, Nika Teharani, Alireza Karimi for performing data analysis and CBDepot.eu for sharing the CBD at shipping costs. Experiments were funded by an NWO VIDI grant to LG.

## Contributions

AAZ, MMC, A.R. analyzed the data, A.S. and L.G. performed the experiments, P.Ö, K.A, T.A. performed specific analyses, L.G. designed the experiments and wrote the first draft of the article together with AAZ and AS, AAZ, MMC, A.R. and A.S. edited the draft.

## Cannabidiol decreases CA3 but not MEC input to CA1 ripples – Supplement

**Fig. S1.**
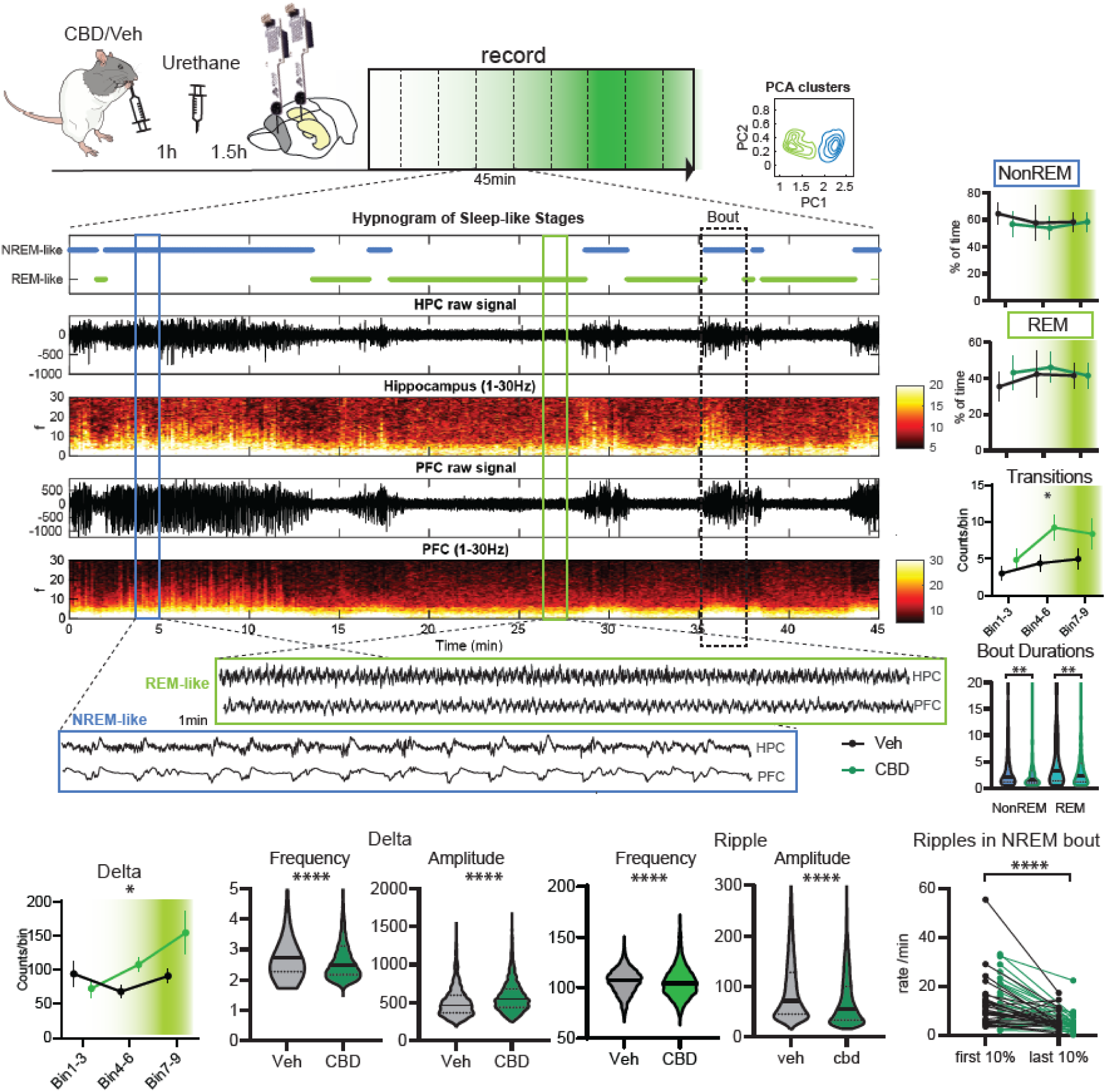
Sleep-like states: Animals received either oral CBD or vehicle and 1h later an urethane injection (Fig. 1A). Urethane anesthesia is known to display NREM- and REM-like states that contain the usual microstructural events such as slow oscillations (0-1.5Hz), delta waves (0-4Hz), ripples, and theta (4-8Hz), as can also be observed in our data (Fig.1B). With a principle component analysis we identified the two sleep-like states (Fig. 1B), splitting our recordings into NREM-like and REM-like periods. CBD induced more state transitions and deepening of NREM with more, slower and larger delta waves in this state (Fig. 1), corresponding to the previously reported extension of NREM seen in natural sleep recordings ^1^. We could also replicate that under CBD ripples during NREM-like state were slower and smaller but similar to ripples in natural sleep generally occurred more in the beginning of each NREM bout than the end. Thus, we replicate our previous CBD findings reported from natural sleep in sleep-like states as well as providing evidence for sleep-like dynamics of our anesthesia recordings, confirming the validity of the model.

**Fig. S2.**
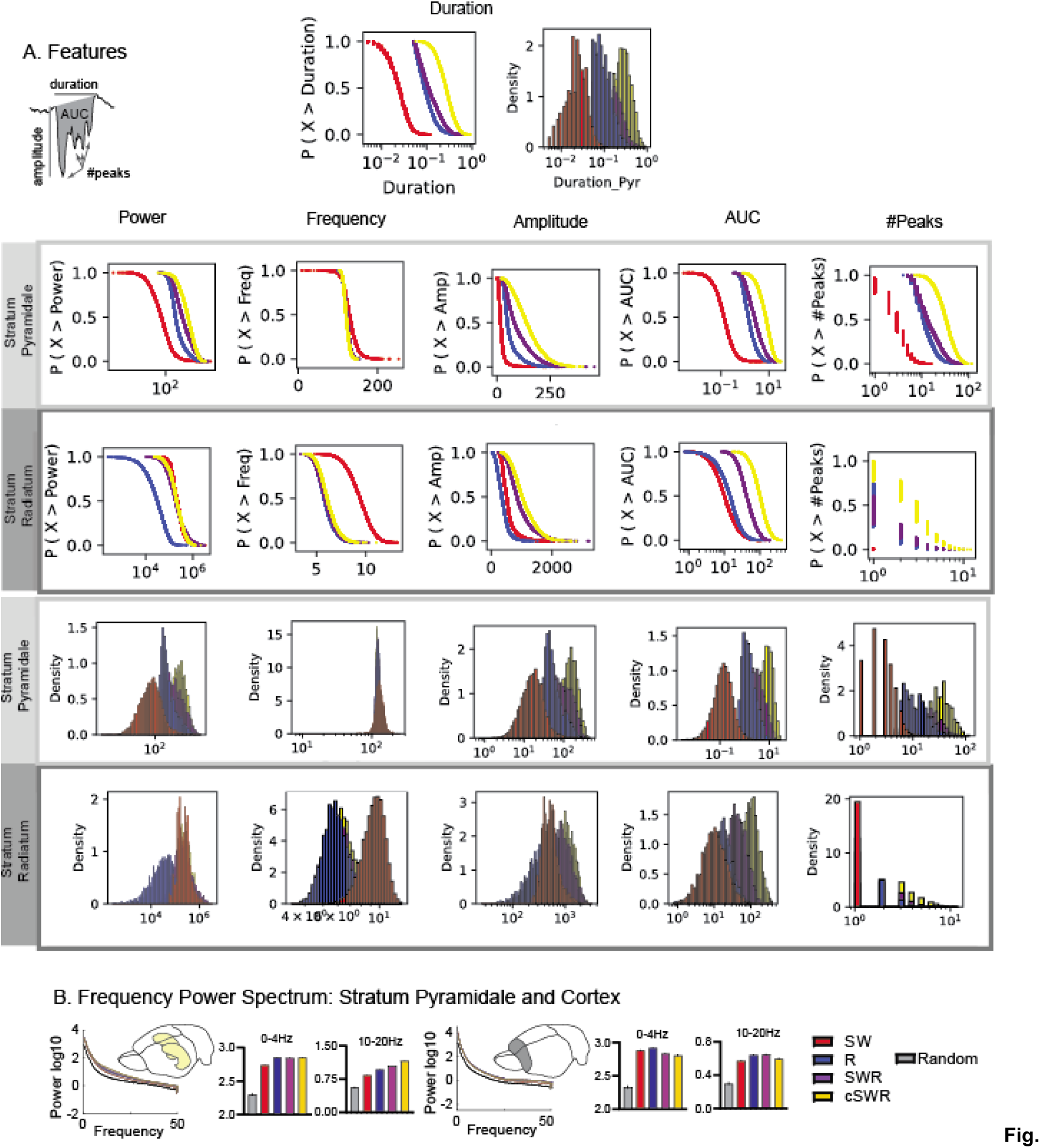
Event Characteristics: A. Shown are the features for the four types of events for both stratum pyramidale and radiatum. B. Power spectrum for stratum pyramidale of hippocampus and for the prefrontal cortex (prelimbic electrodes site) for delta and spindle range.

**Fig. S3.**
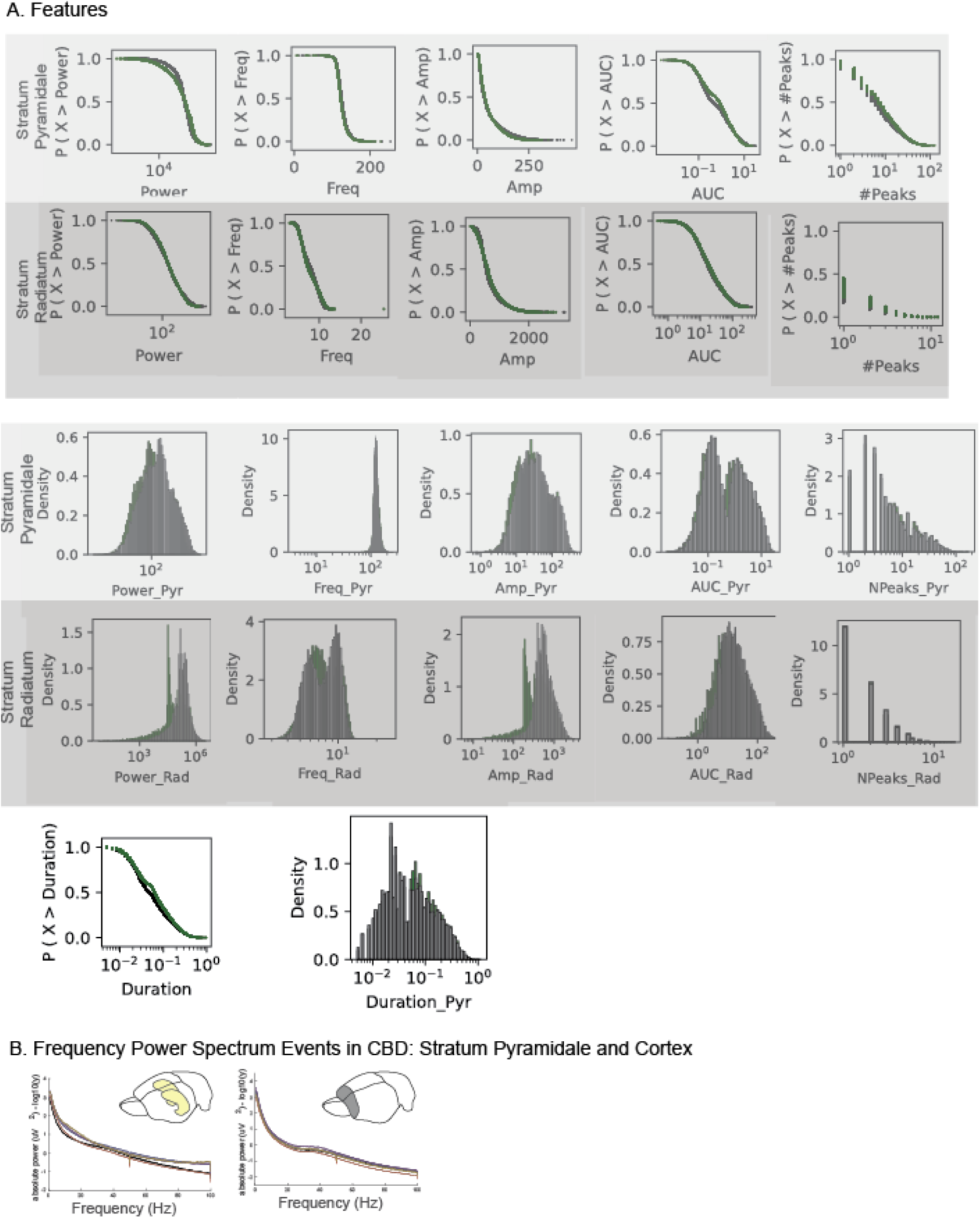
Event Characteristics CBD: A. Shown are the features for the four types of events for both stratum pyramidale and radiatum. B. Power spectrum for stratum pyramidale of hippocampus and for the prefrontal cortex (prelimbic electrodes site).

**Fig. S4.**
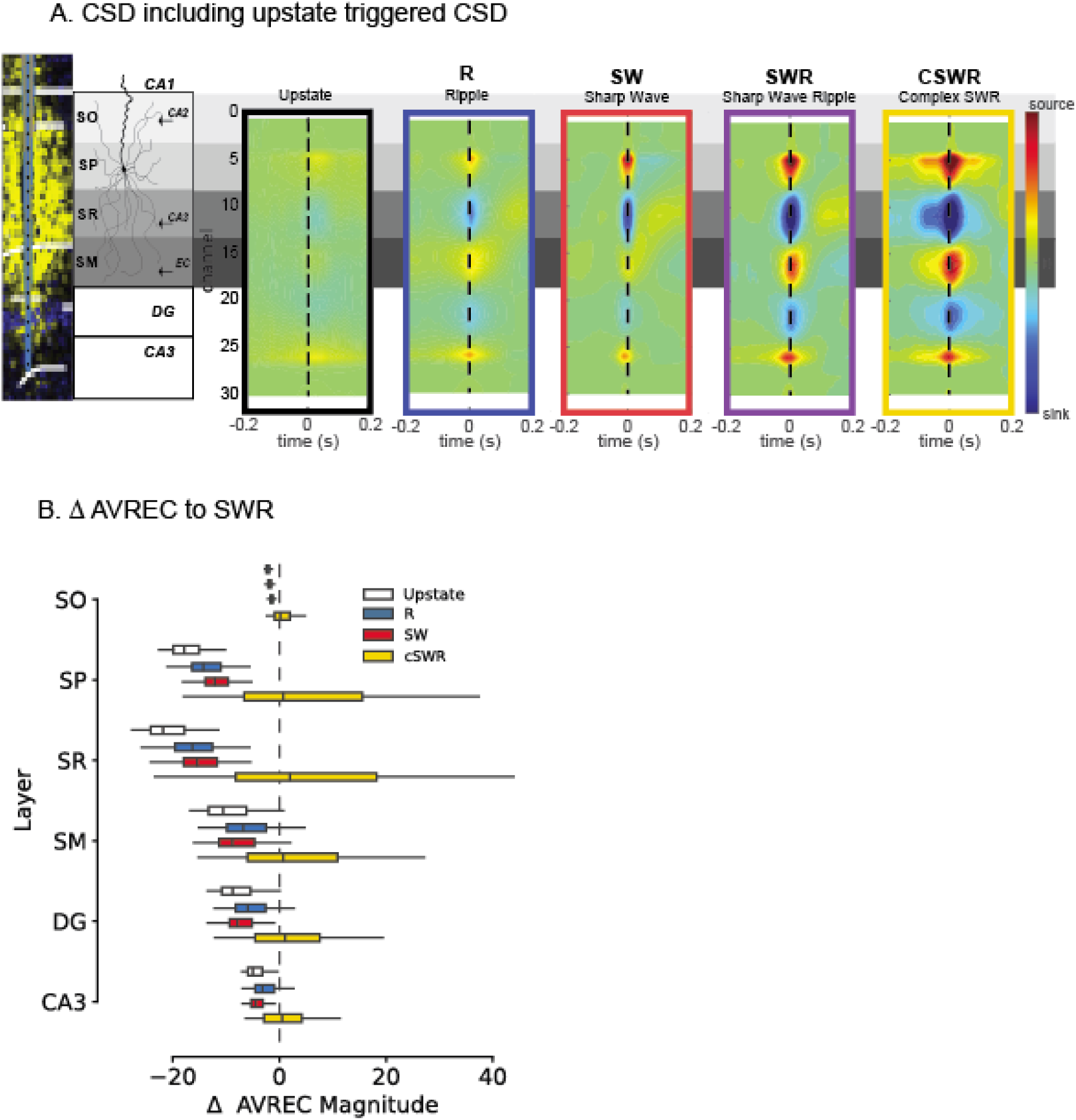
Upstate CSD analysis: A. Shown are the CSD maps for the types and one triggered on the upstate. B. AVREC relative to the SWR events for the other types and upstate.

